# CAncer bioMarker Prediction Pipeline (CAMPP) - A standardised and user-friendly framework for the analysis of quantitative biological data

**DOI:** 10.1101/608422

**Authors:** Thilde Terkelsen, Anders Krogh, Elena Papaleo

## Abstract

**Motivation:** Recent improvements in -omics and next-generation sequencing (NGS) technologies, and the lowered costs associated with generating these types of data, have made the analysis of high-throughput datasets standard, both for forming and testing biomedical hypotheses. Alongside new wet-lab methodologies, our knowledge of how to normalise bio-data has grown extensively. By removing latent undesirable variances, we obtain standardised datasets, which can be more easily compared between studies. These advancements mean that non-experts in bioinformatics are now faced with the challenge of performing computational data analysis, pre-processing and visualisation. One example could be the analysis of biological data to pinpoint disease-related biomarkers for experimental validation. In this case, bio-researchers will desire an easy and standardised way of analysing high-throughput datasets.

**Results:** Here we present the CAncer bioMarker Prediction Pipeline (CAMPP), an open-source R-based wrapper intended to aid non-experts in bioinformatics with data analyses. CAMPP is called from a terminal command line and is supported by a user-friendly manual. The pipeline may be run on a local computer and requires little or no knowledge of programming. CAMPP performs missing value imputation and normalisation followed by (I) k-means clustering, (II) differential expression/abundance analysis, (III) elastic-net regression, (IV) correlation and co-expression network analyses, (V) survival analysis and (IV) protein-protein/miRNA-gene interaction networks. The pipeline returns tabular files and graphical representations of the results. We hope that CAMPP will assist biomedical researchers in the analysis of quantitative biological data, whilst ensuring an appropriate biostatistical framework.

**Availability and Implementation:** CAMPP is available at https://github.com/ELELAB/CAMPP

## Introduction

Utilisation of sensitive and specific biomarkers for disease diagnosis, prognosis and monitoring, is an attractive alternative to many of the current methods in use. The presence and levels of certain tissue-derived molecular markers can help distinguish subtypes in heterogeneous diseases such as cancer (Dai, et al., 2016; Duffy, et al., 2017). Biomarkers may also be predictive of patient outcome and responsiveness to treatment (La Thangue and Kerr, 2011; Vieira and Schmitt, 2018). Moreover, levels and composition of circulating blood biomarkers seem ideal for non-invasive patient screening both before and after the onset of disease (Tang, et al., 2017; Uttley, et al., 2016). Alas, pinpointing robust cancer biomarkers may be a difficult endeavour and in a review from 2014 Yotsukura and Mamitsuka (Yotsukura and Mamitsuka, 2015) showed that out of 7720 publications on biomarkers usage (see original publication for search criteria), only 407 of these were patented and none had obtained FDA approval. One of the main limitations of biomarker research often relates to small sample size, yielding over-fitted and unreproducible results (Berghuis, et al., 2017; Kern, 2012; Tiberio, et al., 2015; Yotsukura and Mamitsuka, 2015). Other pitfalls include lack of standardised data curation (Ioannidis, et al., 2009), inappropriate statistical analysis and lack of validation (Berghuis, et al., 2017; Kern, 2012; Nicolle, et al., 2019). Evaluation of marker specificity and sensitivity is pivotal as most cancer biomarkers have high false positive rates due to the fact that a range of non-cancerous events may cause changes in levels of specific biomolecules. Blood-based biomarkers are especially dynamic and can be affected by time of sampling, patient diet, medication and other biological variances which are extremely difficult to take into account (Tiberio, et al., 2015; Yotsukura and Mamitsuka, 2015). Despite these limitations, molecular markers such as mRNA, proteins, lipids and small RNAs remain a promising strategy for diagnosis and subtyping of cancer or other chronic diseases. With the advancements in the field of high-throughput data and the increased attention to the issues of data normalization and statistical modelling, some drawbacks of biomarker mining may be overcome in a foreseeable future (Minciacchi, et al., 2017).

Central to the identification of novel disease markers is the bioinformatic analysis of high-throughput biological data (Alcaraz, et al., 2017; Ghosh and Poisson, 2009; Merrick, et al., 2011; Papaleo, et al., 2017; Wang, et al., 2017). By applying different statistical tests and machine learning approaches, researchers can go from large datasets with quantitative measurements to a few biomolecules of interest (Huang, et al., 2016; Kursa, 2014; Malta, et al., 2018; Soneson and Delorenzi, 2013; Swan, et al., 2013). The potentials of these markers may then be validated experimentally and subsequently supported by independent patient cohorts and/or through clinical trials. Ideally, biomarker studies of the same disease should be reasonably comparable, however, discrepancies of results are not at all uncommon. While some of these differences arise from variances in study design and experimental procedures, a significant proportion is due to alternating and sometimes inappropriate data normalisation and bioinformatic analysis pipeline (McDermott, et al., 2013; Qin and Levine, 2016; Siu, et al., 2016). Standardising the framework for detection of biomarker candidates both in the wet lab (Khan, et al., 2017), as well as the dry lab, should enable researchers to more directly compare results across different studies and starting materials (Siu, et al., 2016; Witwer, 2015).

Here we present the CAncer bioMarker Prediction Pipeline (CAMPP), an R-based command-line wrapper for the analysis of high-throughput data. The intention behind CAMPP is to provide non-experts in computational biology and bioinformatic software-users with an easy, quick and standardised way of screening for potential disease markers, and other biomolecules of interest, prior to experimental validation. CAMPP may be applied to a variety of quantitative biological data, here among; expression data and mass-spectrometry data. CAMPP may be used to perform different data normalizations and bioinformatic/biostatistical analysis. Although distributional checks etc., are automatically implemented in CAMPP, it is vital that users have a good understanding of the generation and post-processing of their input data, to avoid gross statistical violations and misinterpretation of results. Generally, we have aimed for a pipeline which is intuitive and well-documented, as is good practice (List, et al., 2017).

### System

#### Accessibility

The CAncer bioMarker Prediction Pipeline (CAMPP) is open source and may be downloaded from the github repository: https://github.com/ELELAB/CAMPP

The repository contains R-scripts, data examples, as well as a user manual. The pipeline consists of three R-scripts. (I) CAMPPInstall.R, (II) CAMPPFunctions.R and (III) CAMPP.R.

#### Requirements

CAMPP is built on a variety of R-packages and as such the user must have installed R and preferably Rstudio. Furthermore, Macbook users must have Xcode installed (https://developer.apple.com/xcode/), while windows users must ensure they have some equivalent of command line tools.

Specifics on R-packages required (versions and how to troubleshoot) is specified in the user manual: https://github.com/ELELAB/CAMPP/blob/master/CAMPPManual.pdf

Currently CAMPP is a command-line tool run using Rscript, which is automatically acquired when installing R and Rstudio (most likely path; /usr/local/bin/Rscript). Minimum requirements; R version 3.5.1 and Rstudio version 1.1.463.

#### Set-up

The CAMPPInstall.R only needs to be run the first time the pipeline is used, as this script ensures that all required R-packages will be installed correctly. The CAMPPFunctions.R contains custom functions sourced by the CAMPP.R script (main script) and as such these two scripts must be located in the same working directory (folder) in order for the pipeline to run.

## Methods and Implementation

The CAncer bioMarker Prediction Pipeline may be used to perform a variety of analysis, including preliminary data management, summarized in Table 1 and Figure 1A below. As analysis with CAMPP is standardised, the pipeline accepts most biological datasets with expression/abundance values from microarrays, mass spectrometry and high-throughput sequencing platforms. For clarity, we will in brief go over some of the assumptions underlying data input and subsequent analysis, along with mandatory parameters.

**Table 1.**
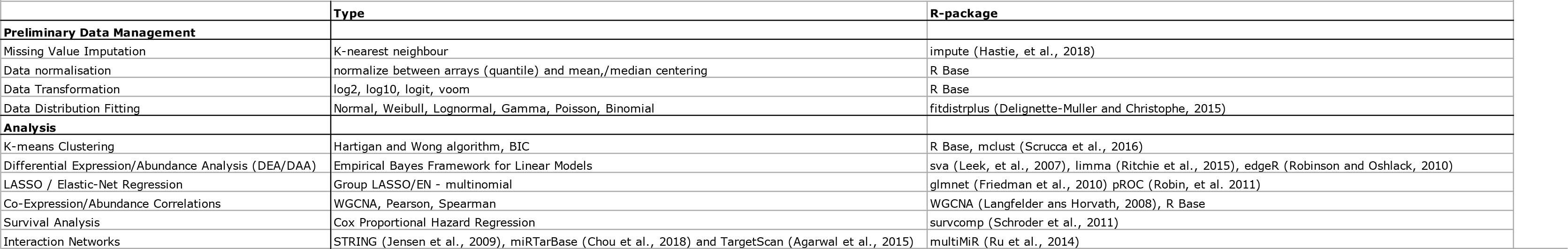
Table summarizing preliminary data management and analyses implemented in CAMPP, along with specific methods and underlying R-packages.

**Figure 1:**
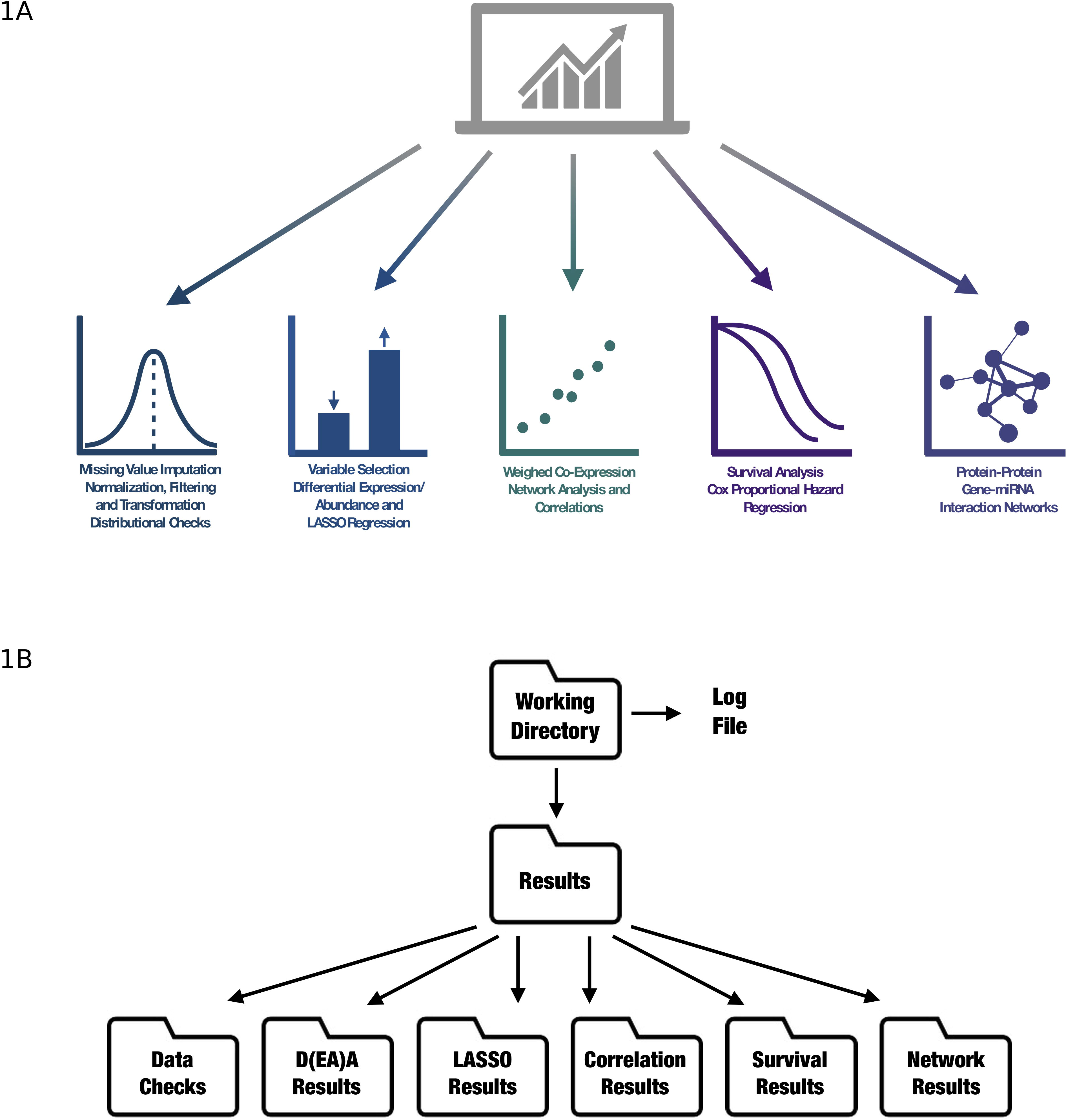
The diagram depicts the methods employed by the CAncer bioMarker Prediction Pipeline (CAMPP).

### User Input

As a minimum the user must provide; (I) a matrix with expression/ abundance values (.xlsx or .txt file), with variables as rows and samples as columns, (II) a metadata file (.xlsx or .txt) which should contain at least two columns, a column with sample ids, matching the column names in the expression matrix and a column specifying which group (disease state, treatment etc.) a given sample belongs to. Lastly, the user has to specify which type (variant) of data is provided for analysis, current options are; *array, seq, ms* or *other*.

### Value Imputation and Pseudocounts

If the input data contains missing expression values, these will automatically be imputed using local least squares imputation, LLSI (spearman correlation) (Stacklies, et al., 2007) or k-nearest neighbour imputation, KNNI (euclidean metric) (Hastie, et al., 2018). Default method is LLSI, as this type of imputation has been shown to perform better on expression data (Celton, et al., 2010). However, for datasets with many missing values, or if the LLS fails, KNN is applied instead. If the user wants to log-transform the data before analysis and the dataset contains zeros, a pseudocount will be added to enable transformation.

### Data Normalization, Filtering and Transformation

Based on what data variant is input (array, seq, ms), different steps of normalization will be performed. For RNA sequencing data (seq) variables with low counts over all groups (tissue, treatment) are filtered out, library sizes are scaled (normalization method is weighted trimmed mean of M-values, TMM) (Robinson and Oshlack, 2010) and data are *voom* transformed (Soneson and Delorenzi, 2013). For microarray data, (array) data are log2 transformed and either quantile normalised (normalizeBetweenArrays) (Bolstad, et al., 2003) or standardised using mean or median (Luo, et al., 2010). For mass spectrometry data or if data variant other is specified (ms or other), data are log transformed (log2, log10 or logit, argument −t) and standardised using mean or median (argument −z) (Luo, et al., 2010).

It should be noted that CAMPP does not perform within-array-normalization, standard for two-color array data (Quackenbush, 2002). Within-array-normalization would be complex to standardise, taking into account array type, spots per print-tip group, array weights etc. As this is beyond the scope of CAMPP, intended for downstream data analysis, within-array-normalization must have been done before hand (see *limma* manual for more information (Ritchie, et al., 2015)).

### Preliminary Data Distributional Checks

As a default the pipeline utilised the R-package *fitdistrplus* (Delignette-Muller and Christophe, 2015) to generate skewness-kurtosis plots (cullen and frey graphs) for n randomly selected variables (genes, proteins etc.), from the input dataset. Distributions are fitted to data by maximum likelihood and parameters of the distribution are estimated with bootstrap-resampling in order to simulate variability (Delignette-Muller and Christophe, 2015). In addition to the cullen and frey graphs, histograms, quantile-quantile and probability-probability plots are returned. In this way the user may evaluate the distributional properties of the data before analysis.

### Clustering of Samples

Before variable selection, the pipeline may be used to perform k-means clustering analysis. CAMPP will test a number of centroids, the exact number of which will depend on the size of the dataset. After clustering CAMPP employs the R-package *mclust* (Scrucca, et al., 2016) to evaluate which number of clusters is “optimal” for the input data, based on the bayesian information criterion (BIC), e.g. a comparison of maximum log likelihoods with a penalty for overfitting subtracted. K-means clustering with CAMPP will return a multidimensional scaling (MDS) plot depicting the “best” clustering of samples. The flag denoting k-means clustering, takes a string specifying which column from the metadata file should be added as labels to the plot(s), allowing the user to compare the results of clustering with the available sample information. In addition to MDS plots, the pipeline returns the original metadata file with one or more columns appended specifying which cluster which sample was assigned to. If desired, the user may re-run CAMPP, using the k-means column for variable selection.

### Variable Selection with Differential Expression Analysis and Elastic-Net Regression

Essentially, CAMPP should provide biomedical researchers with a tool they can use to perform standardised “computational screens” of high-throughput data, with the purpose of selecting a small subset of interesting biomolecules. The selected candidates, with potentials as diagnostic or prognostic disease biomarkers, may then be studied and validated in a laboratory setting. For the purpose of variable selection CAMPP employs *limma* (linear models for microarray data) for differential expression/abundance analysis (DEA, DAA) (Ritchie, et al., 2015). Although *limma* was originally designed for analysis of microarray data and subsequently revised to handle RNA sequencing data, this software is very flexible and has recently been shown to perform very well with quantitative mass spectrometry data (Kammers, et al., 2015; van Ooijen, et al., 2018). In addition to being versatile, *limma* has been shown to work exceptionally well on datasets with small sample sizes (Seyednasrollah, et al., 2015; Soneson and Delorenzi, 2013). This is due to the fact that this software employs parametric empirical Bayes models, which allow for information borrowing across variables, and as a consequence, scaling of residual variances (Morris, 1983; Ritchie, et al., 2015). Lastly, *limma* is relatively fast when compared to other methods for DEA (Seyednasrollah, et al., 2015). Differential expression analysis may be performed with correction for experimental batches and/or other confounders, such as tumour immune infiltration scores.

CAMPP also performs least absolute shrinkage and selection operator (LASSO) or Elastic Net (EN) regression with *glmnet* (Friedman, et al., 2010). DEA analysis is automatically performed by CAMPP, whereas the user must specify if EN/LASSO is desired (argument −l). If argument −l is set to 1.0, LASSO regression will be performed, while a value > 0 but < 1.0 will result in Elastic Net (default is 0.5), ridge-regression is not supported.

EN/LASSO may be performed in two ways, (I) the dataset is split into training and testing subsets, k-fold cross validation is performed on the training dataset, followed by estimation of specificity and sensitivity (area under the curve = AUC, (Robin, et al., 2011)) using the test dataset, or (Il) k-fold cross validation is performed using the full dataset, no AUC is reported. The pipeline will automatically estimate whether the input dataset is large enough to split into training and test subsets and whether EN/LASSO is advisable to perform altogether. CAMPP will perform regression analysis 10 times and output bar-plots of cross-validation errors and AUCs for each run. In addition to a tabular .txt file with the results of variable selection (consensus of 10 runs), a file with the overlap between DEA and LASSO/EN will be generated.

### Weighed Co-Expression Network Analysis and Correlation Analysis

CAMPP may be used to perform pairwise correlation analysis (Spearman or Pearson), with testing for correlation significance and correction for multiple testing (FDR). The two datasets provided do not need to have the same dimensions, but these must have a partial overlap of variables and sample IDs. Filtering, normalization, batch-correction and transformation may be performed as described above. Results of correlation analysis will be summarized in a plot with correlation scores of all variables tested, as well as in individual correlation plots of the variables which remain significant after correction for multiple testing.

The user may perform Weighted Co-Expression Network Analysis. For this type of analysis CAMPP relies on the R-package *WGCNA* (Langfelder and Horvath, 2008). To reduce the contribution from low correlations, mainly assumed to be noise, the WGCNA software estimates a soft thresholding powers for exponentiation and returns a plot of these. If this plot shows that the scale-free topology fit does not reach a value > 0.8 for any tested power, then the dataset may encompass either a large biological heterogeneity or technical variability. If experimental batches are known these should be corrected for and co-expression analysis re-run, either way, the user should consider whether WGCNA is suitable for the dataset in question. Co-expression analysis with CAMPP (*WGCNA* (Langfelder and Horvath, 2008)) will result in a plot of variable clustering, before merging and after merging of modules (modules which have less than 25% dissimilarity will be merged by default). A heatmap showing strength of variable co-expression within each module will be generated, if the module contains <= 100 variables - more than this will yield an unreadable plot. In addition, CAMPP will return the top most interconnected variables from each module with accompanying interconnectivity score plots.

### Survival Analysis - Pinpointing Prognostic Biomarkers

With CAMPP users may perform survival analysis with cox proportional hazard regression with *survcomp* (Schroder, et al., 2011). To run survival analysis the provided metadata file must contain at least four columns; age = age in years at diagnosis, surgery or entry into trail, outcome.time = time until end of follow-up, censuring or death in weeks, months or years, outcome = numeric vector specifying censuring = 0 or death = 1 and survival = numeric vector specifying if survival information exist, no info = 0 and info available = 1. If the user wishes to correct for potential confounders other than age (e.g. tumour grade, drug-treatment etc.) these should also be included in the metadata file. Confounders should be provided as a list of commas separated names, which match the names of columns in the metadata file.

The CAMPP pipeline checks two underlying assumptions of the cox model before performing survival analysis (I) a linear relationship of continuous covariates with log hazards and (II) proportional hazards of categorical and continuous covariates, e.i. constant relative hazard (Liu, et al., 2018). If the requirement of linearity is not fulfilled, cubic splines will be added to the covariate(s) in question.

### Interaction Networks

After variable selection, the user may generate protein-protein and/or miRNA-gene interaction networks. If gene expression data are used as input for CAMPP, protein-protein interactions are retrieved from the *STRING database* (Jensen, et al., 2009) and pairs where both genes (proteins) are differentially expressed are extracted. Low scoring interactions are filtered out, here defined as < 25th quantile of all scores in the database. The pipeline can accept a range of gene identifiers, including gene symbol, gene identifiers, transcript identifiers etc. If miRNA expression data are used as input, then miRNA-gene interaction pairs are retrieved from either *miRTarBase* (validated targets) (Chou, et al., 2018), *TargetScan* (predicted) (Agarwal, et al., 2015) or a combination of both (Ru, et al., 2014), as specified by the user. Low scoring predicted interactions from TargetScan are filtered out, low scoring defined as < 25th quantile of all scores in the database. Mature miRNA identifiers or miRNA accession are allowed as input. Because only differentially expressed miRNAs are available for analysis, nothing is known about the paired genes, e.g. these are reported without statistics. If the user has both gene and miRNA expression values from the same sample cohort, both protein-protein and miRNA-gene pairs are retrieved and the results are combined. In this case, expression directionality of both miRNAs and genes are known, and the pipeline will therefore only return pairs where the fold changes of gene and miRNA are inverse, one up-regulated and the other down-regulated. Interactions network analysis with CAMPP will result in a tabular .txt file with all extracted interactions, including logFCs, FDRs and interaction scores. This file may be used for visualization of networks with *cytoscape* (Shannon, et al., 2003) or another similar tool. In addition, a plot of the top 100 “best” interactions (based on absolute logFCs and scores) are returned. One file and one plot will be generated for each pairwise group contrast which returned differentially expressed genes.

### Output

More generally, CAMPP.R results are returned as tabular .txt files and .pdf graphics files. Figure 1B shows the structure of the CAMPP output with all flags specified.

#### Case Study 1

##### Analysis of Single-Channel Microarray Data - Variable Selection, WGCNA and Gene-Gene Interactions

For the testing of an array dataset, we used mRNA expression data from single-channel microarrays (Jabeen, et al., 2019). The dataset contained 80 breast tumour samples with expression quantified for ~ 15.000 mRNAs. Breast tumour samples were stratified into subtypes as follow; 38 Luminal A, 11 Luminal B, 12 Luminal B Her2 enriched, 9 Her2 enriched and 10 triple negative breast cancer (TNBC). Clinical data contained, among other things, information on experimental batch (array), tumour grade, tumour infiltrating lymphocyte scores, hormone receptor statuses and patient outcome. Data were background-corrected for ambient intensities before analysis. The dataset contained a small number of missing values between 0-3% per row.

We used CAMPP to perform variable selection, e.g. differential expression analysis and elastic net regression. For DEA cut-offs for log fold change and adjusted p-values (FDR) were set to 1 and 0.05, respectively. Missing values were imputed, data were log2 transformed and normalised between arrays. Data were corrected for experimental batches, specified as a column in the metadata, by setting the flag specifying batch-correction. In addition, the immune score status was added to the design matrix as a covariate.

Variables with the ability to partition patient estrogen receptor status (ER+ vs ER−) were selected with elastic-net (alpha = 0.5) and DEA. For this contrast patient distribution across the two groups was balanced with enough samples to divide the data into training and testing sets (automatically done by CAMPP). As another example, BC subtypes were compared, however, as the group of Her2-enriched tumours only contained 9 samples, elastic-net regression (alpha 0.5) with CAMPP was performed without splitting the dataset into a training and testing set, e.g. all data were used for training with 10-fold cross-validation. As the lack of a test set increases the chance of overfitting significantly, elastic-net results were not accessed individually but simply provided support for DEA results.

Figure 2 shows an example of the data checks performed with *fitdistrplus* (Delignette-Muller and Christophe, 2015) on a set of n (default is 10) randomly extracted variables from det dataset. The gene used as an example in Figure 2 is FAM27E2 (Family With Sequence Similarity 27 Member E2), randomly selected from the ten data check plots. Distributional checks for the other nine genes may be found in the zip folder SupplementaryCSupS1.zip.

**Figure 2:**
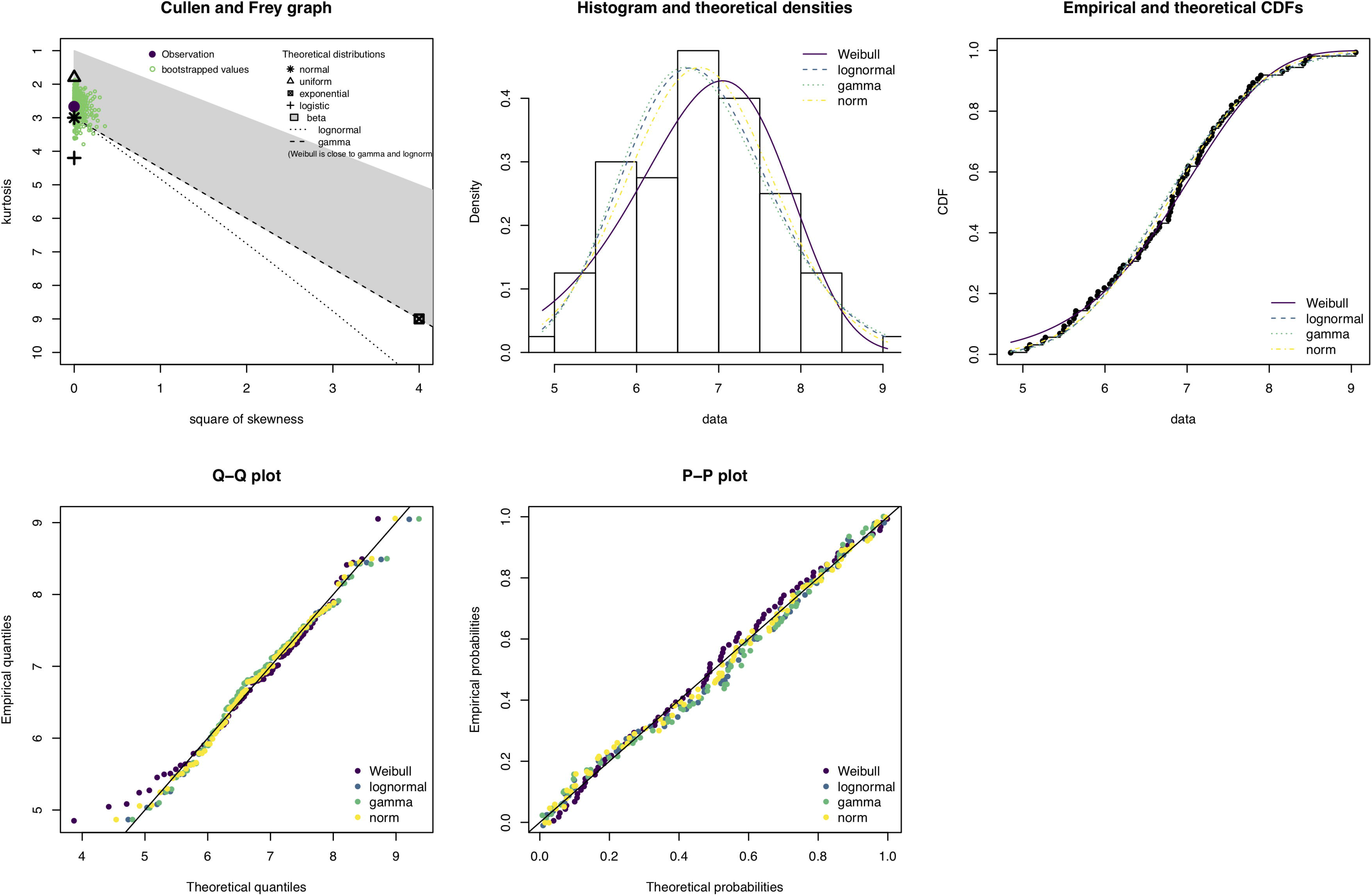
Structure of the output files and directories produced by a CAMPP run.

Figure 3 shows the number of up and down regulated genes from DEA overlapped with results of elastic-net regression, for the comparison of estrogen receptor status (ER− vs ER+). Figure 3 also contains a MDS plot for data-overview (corrected for experimental batch (Leek and Storey, 2007)) and statistics on cross-validation error and area under the curve (AUCs) for each of the 10 elastic-net runs. Lists of DE genes, results of LASSO regression and all plots may be found in the zip folder SupplementaryCS1.zip (see Github repository).

**Figure 3:**
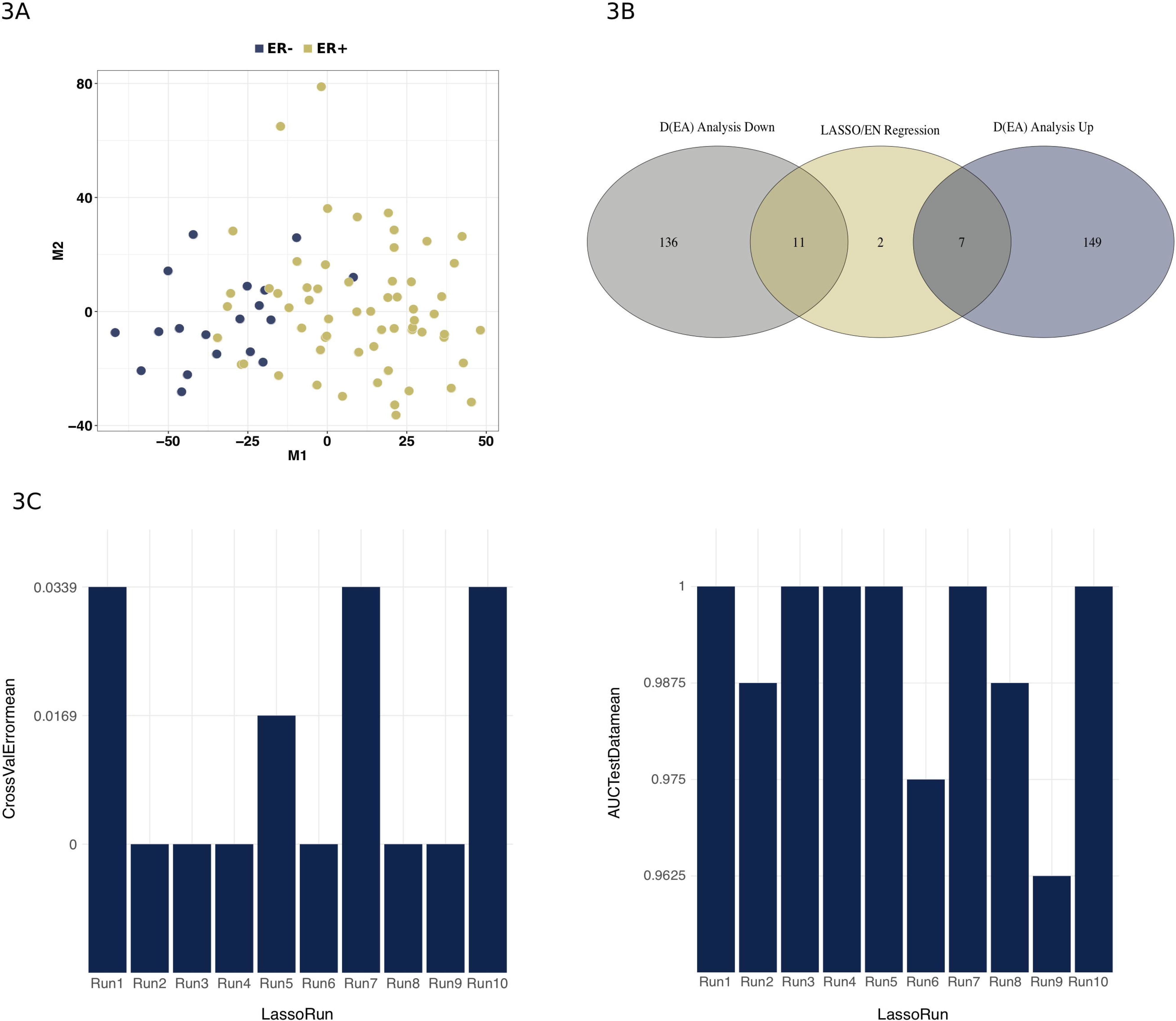
Results of gene selection using differential expression analysis and elastic-net regression. The dataset contained ~ 15.000 genes and 80 samples, groups used for contrast were estrogen positive (n = 61) vs estrogen negative samples (n = 19). Figure 3A is a multidimensional scaling plot showing the partitioning of samples (based on all genes), colored by estrogen status. Figure 3B shows the overlap of results from elastic-net regression (alpha = 0.5) and differential expression analysis with significance cutoffs logFC > 1 or < −1 and FDR < 0.05. Figure 3C depicts the performance statistics for elastic-net regression, e.g. 10-fold cross validation errors and area under the curve (AUC) scores for test set. Elastic-net is run 10 times with different random seeds.

As seen from Figure 3B a total of 147 genes were found to be down-regulated in ER− vs ER+ (Up in ER+), while 156 genes were up-regulated in ER− vs ER+ (down in ER+). Elastic-Net regression resulted in 20 genes, out of which nine overlapped with DEA results. Plots of cross-validation errors and AUCs displayed nice convergence, with AUCs ranging between 0.96-1.0, e.g. high sensitivity and specificity for the test set. A quick look into the nine genes from the overlap between DEA and elastic-net, revealed that two of these were Estrogen Receptor 1 and RAS Like Estrogen Regulated Growth Inhibitor, both up-regulated in the ER+ samples compared to ER− samples, in accordance with expectation (Habashy, et al., 2011). Two of the genes which were up-regulated in ER−negative samples encoded for solute carriers, known to be associated with more aggressive types of breast cancer (El-Gebali, et al., 2013; Yen, et al., 2018), supporting their over-expression in the ER-negative vs ER-positive group (Rakha, et al., 2007). The heatmap in Figure 4 shows the partitioning of breast tissue samples based on the Consensus set of variables from DE analysis and Elastic-net regression.

Results of the DEA and elastic net regression with BC subtypes, may be found in the SupplementaryCS1.zip folder in the Github repository. In summary, a total of 290 genes were differentially expressed between subtypes, out of which 20 were also identified by elastic-net regression - in total elastic-net returned 42 genes. The set of 20 consensus genes encompassed many well-known genes with the potential to distinguish subtypes. In addition to three pam50 genes; ERBB2, ESR1 and FOXA1 (Parker, et al., 2009), the consensus set included; C1orf64 (ER−related factor, ERRF) which was down-regulated in TNBC, in accordance with literature (Naderi, 2017; Su, et al., 2012), CDK12, which was up-regulated in Her2 samples, supported by literature indicating that a total of 71% of Her2-enriched tumours over-express this gene (Paculova and Kohoutek, 2017). Other genes of interest were CYP2B6 (Lo, et al., 2010), MYLK3 (D’Amato, et al., 2015) and SLURP1 (Gao, et al., 2014).

**Figure 4:**
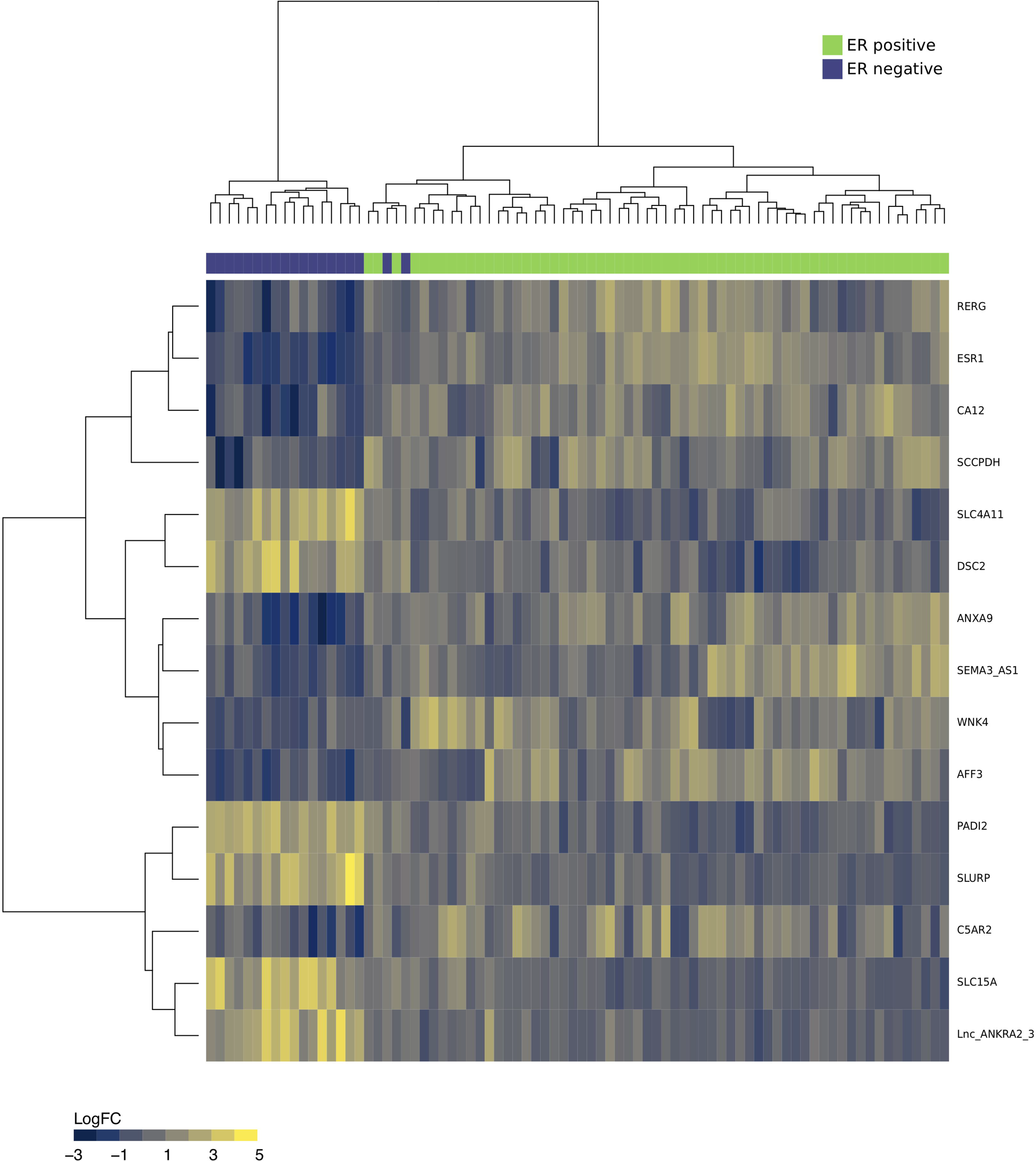
The heatmap in Figure 4 shows the partitioning breast cancer tissues into estrogen receptor positive (ER+) samples and estrogen receptor negative (ER−) samples, based on the consensus set of variables from differential expression analysis and elastic-net regression. Green = ER+ samples and Purple = ER− samples. Color scale of heatmap (blue to yellow) denotes log2 fold change.

As a side note, subtyping of the example dataset, was performed using immuno-histochemistry and not pam50 classification, also, the set only contained about half the pam50 set of genes, e.g. we were not expecting a full overlap of pam50 genes.

In addition to variable selection we used CAMPP to perform Weighted Gene Co-expression Network analysis (WGCNA). In this example, WGCNA was generated only for the differentially expressed genes. Figure 5 shows a subset of results from WGCNA with genes DE between breast cancer subtypes. Figure 5 sub-figures; (I) 5A, a module dendrogram, (II) 5B, an example of a module heatmap showing co-expression of genes in one of the small modules, module 2, and (III) 5C, an example of a module interconnectivity plot, for the top 25% (default setting) most interconnected genes in module 2. The six genes returned as the most interconnected in this module, were strongly associated with Her2-enriched BC subtype. These genes included ERBB2 (HER2 itself), GRB7 (a Pam50 classifier gene) and MIEN1, CDK12, PGAP3 and TCAP, all of which are ERBB2 amplicon passenger genes (Cancer Genome Atlas, 2012; Hansen, et al., 2015; Plautz, et al., 2014). These results highlight the utility of the complementary R-frameworks implemented in the CAMPP wrapper.

**Figure 5:**
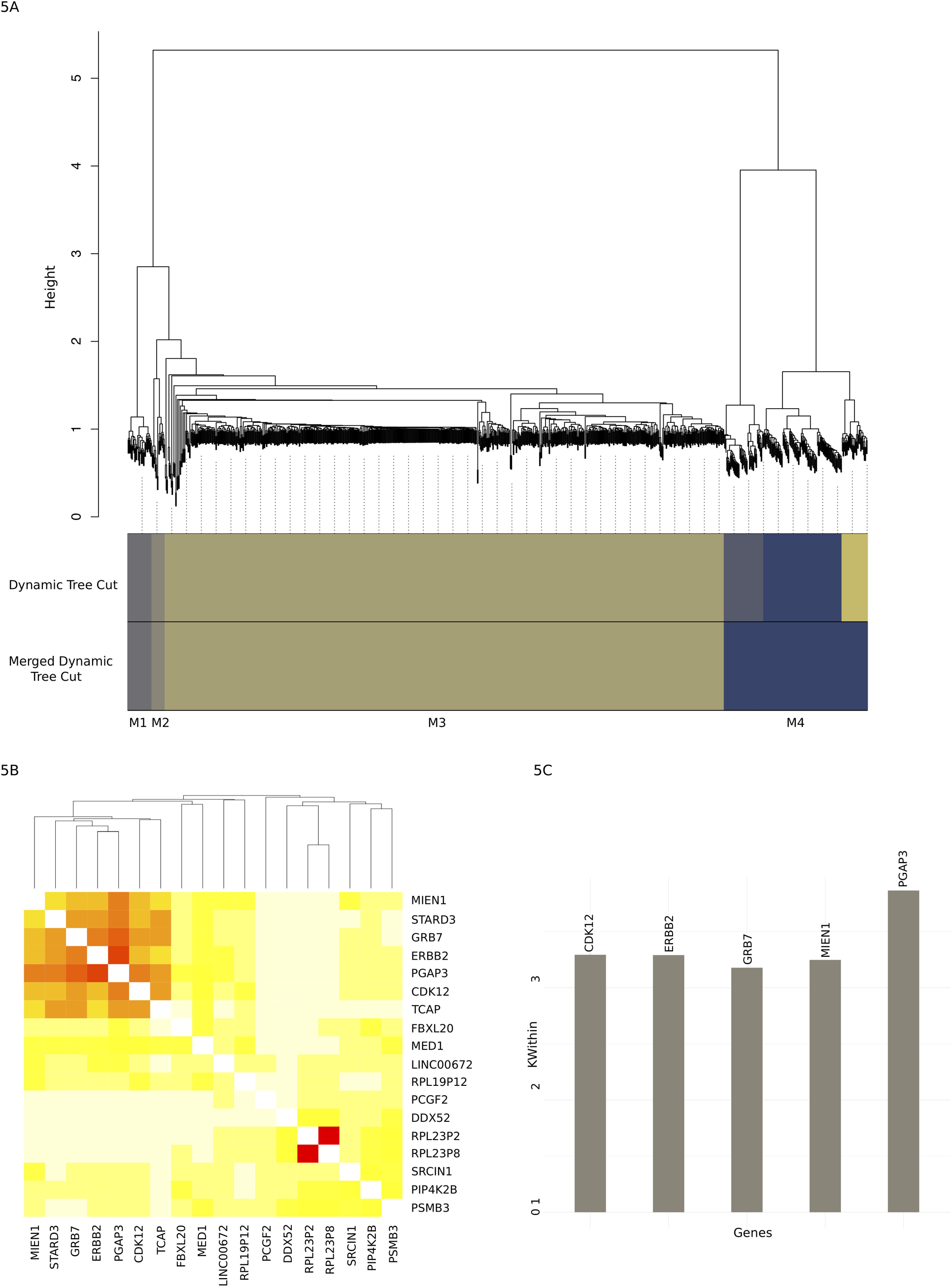
Results of Weighed Gene Co-expression Network Analysis on dataset of ~ 15.000 genes and 80 samples. As the dataset contained more than 5000 variables, WGCNA was performed in a block-wise manner to save computational time, in accordance with the WGCNA reference manual (Langfelder and Horvath, 2008). Figure 5A shows the module clustering tree for the first block as an example. Figure 5B depicts the co-expression heatmap for the small module 2, in which a set of six genes display highly correlated expression patterns. Figure 5C contains the top 25% (in this case five) most interconnected genes from the small module 2, with module interconnectivity scores.

Lastly to evaluate whether the variables found to partition BC subtypes were also known/predicted to interact, we used the CAMPP pipeline to generate protein-protein interaction networks. All differentially expressed genes were included, as log fold changes are useful for this type of analysis. Figure 6 shows the top 100 strongest gene-gene interactions based on absolute logFC and interaction score for the comparison of HER2-enriched vs Luminal A, as an example. Results for comparison of all subtypes pairwise may be found in SupplementaryCS1.zip in the Github repository. From the plot in Figure 6, the two most interconnected genes in the contrast of HER2-enriched vs Luminal A samples, were ERBB2 (HER2) and ESR1 (Estrogen Receptor 1). As expected, ERBB2 was highly up-regulated, while ESR1 was down-regulated in this comparison.

**Figure 6:**
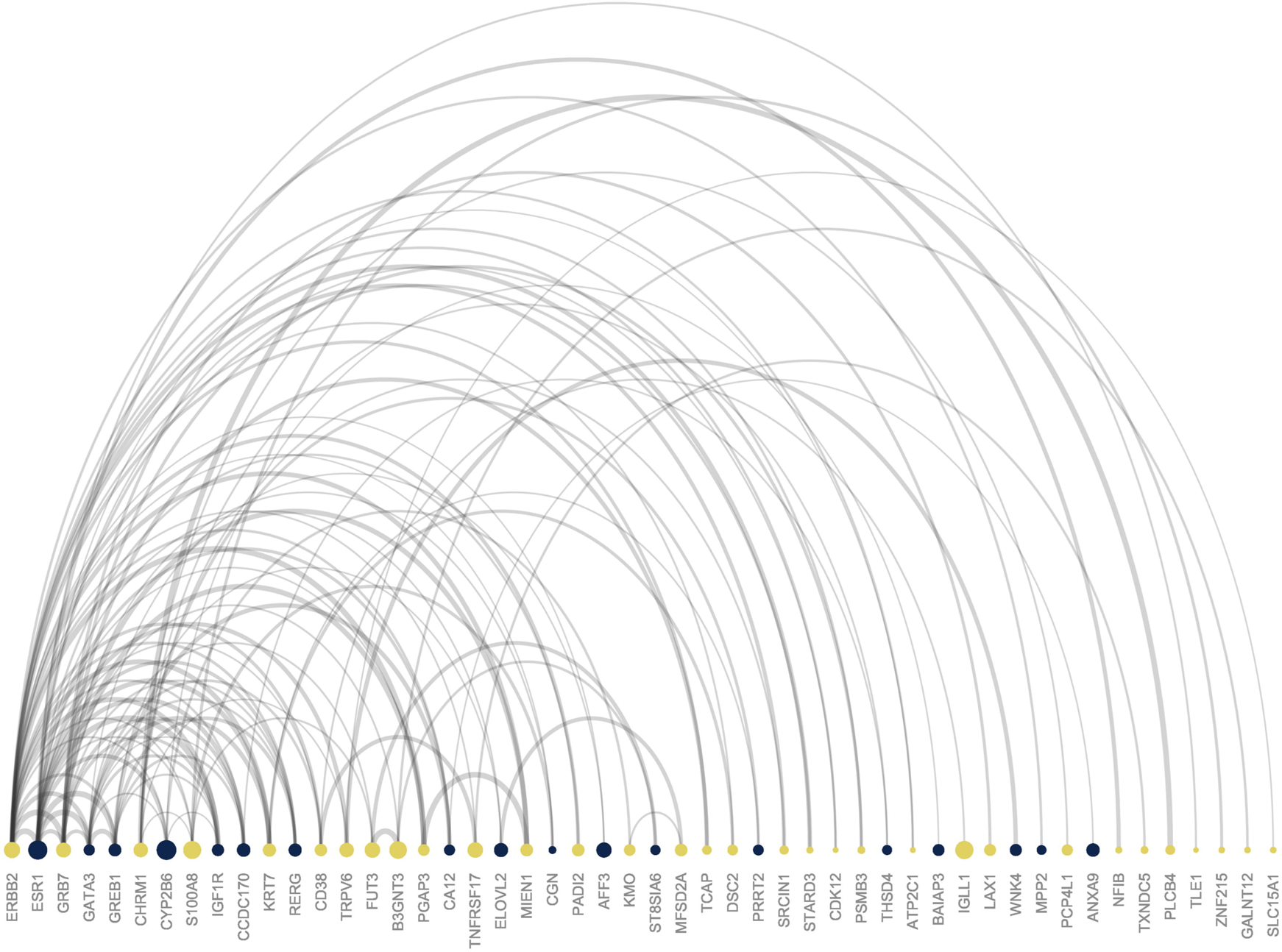
Plot showing the top 100 best protein-protein (gene-gene) interaction pairs from the analysis of HER2-enriched vs Luminal A samples. Colors denote the log fold change of a gene; yellow = up-regulated and blue = down-regulated. The size of the node shows the absolute log fold change, while the ordering from left to right denotes the degree of node interconnectivity. The width of the arch represents the interaction score from the STRING database (Jensen, et al., 2009).

#### Case Study 2

##### Analysis of N-glycans from LC Tandem Mass Spectrometry - Clustering, Tissue-Serum Correlation and Survival Analysis

For case study 2, we analyzed quantitative N-glycan data from liquid-chromatography tandem mass spectrometry (LC-MS/MS). Data were generated from the same cohort of breast cancer samples used for case study 1. However, unlike the mRNA expression data, N-glycans abundances were quantified from tumour and normal interstitial fluids (TIFs and NIFs), not solid tissue. In addition to interstitial fluids we had paired serum samples, enabling us to perform TIF-serum N-glycan abundance correlation analysis with CAMPP. We have published the results of the non-automatized in-depth analysis of these data here; (Terkelsen, et al., 2018). Before differential abundance analysis we used the pipeline to perform K-means clustering of the samples. When running K-means with CAMPP, the user may specify labels to add to the MDS plots generated, in order to see which clinical variables, if any, best explain the observed clusters. Unfortunately for the N-glycans no optimal number of clusters was reported by CAMPP, e.g. the samples could not easily be subdivided into robust clusters. In this case pipeline will output MDS plots for all ks tested (in this case k = 1:10). From evaluation of all k-means plot, the best number of clusters appeared to be two, one corresponding to tumour samples and one to normal samples. Supplementary Figure 1 shows the two clusters with labels.

A clustering analysis we performed differential abundance analysis and elastic-net regression, as described in the section on variable selection and in Case Study 1. No logFC cut-off was set as N-glycan abundances display low variability, cut-off for FDR < 0.05 and for elastic-net regression alpha was set to = 0.5. DAA yielded a total of 20 N-glycan groups (12 up-regulated in cancer vs normal and 8 down-regulated in cancer vs normal), while elastic-net returned six N-glycan groups, all encompassed by the DA set - results may be found in the SupplementaryCS2.zip. Serum correlation analysis and survival analysis were performed on the set of DE N-glycans. Figure 7 shows the results of the correlation analysis with paired N-glycan abundances from TIFs and serum. Three N-glycan groups, GP1, GP37 and GP38 were found to displayed significant correlation scores after correction for multiple testing. Figure 7A shows correlation scores of all tested N-glycans, while 7B shows the individual scatter plots produced for the three significant N-glycan groups. Cox proportional hazard regression was performed with correction for age at diagnosis as well as immune scores (tumour infiltrating lymphocyte scores, TILs), as immune infiltration has been shown to be associated with patient response to treatment and survival (Fridman, et al., 2012). Figure 8 shows the hazard ratios for the DA N-glycan groups with confidence intervals. As seen from the figure, one N-glycan group, GP38, was significantly associated with overall survival, e.g. a high level of TIF GP38 was predictive of a poor patient outcome.

**Figure 7:**
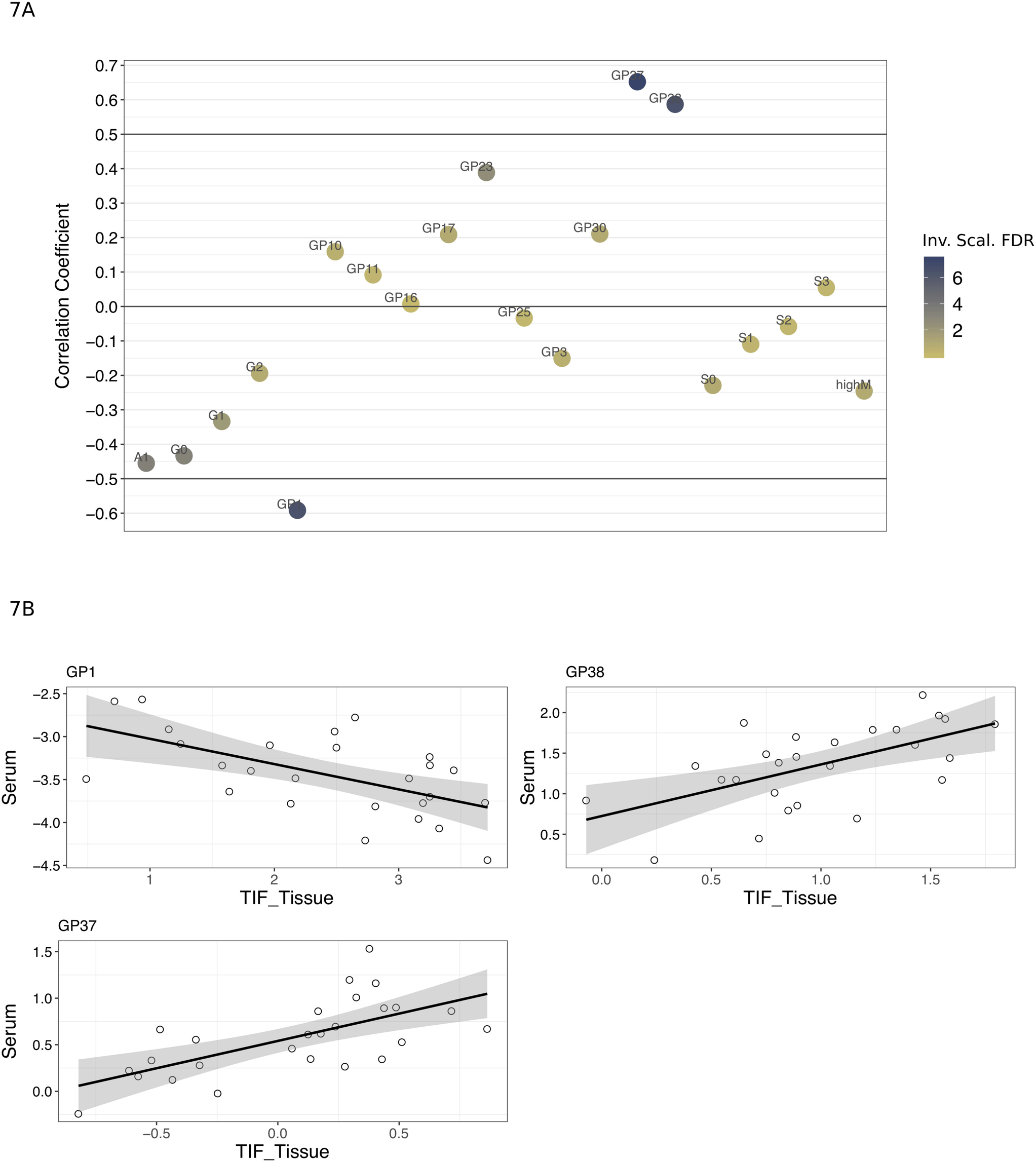
Results of correlation analysis with N-glycan abundances in interstitial fluids and paired serum samples. Dataset contained a total of 103 samples (51 normal interstitial fluids and 52 tumour interstitial fluids) with ~70 N-glycan groups (165 N-glycans). Figure 7A shows the correlation scores for differentially abundant N-glycan groups, three of these, GP1, GP37 and GP38 met the requirement for significance (corr > 0.5 and fdr < 0.05), y-axis = Spearman correlation coefficient. Figure 7B shows the individual correlation plots for the three significant N-glycan groups, x-axis = tumour interstitial fluid abundance and y-axis = serum abundance.

**Figure 8:**
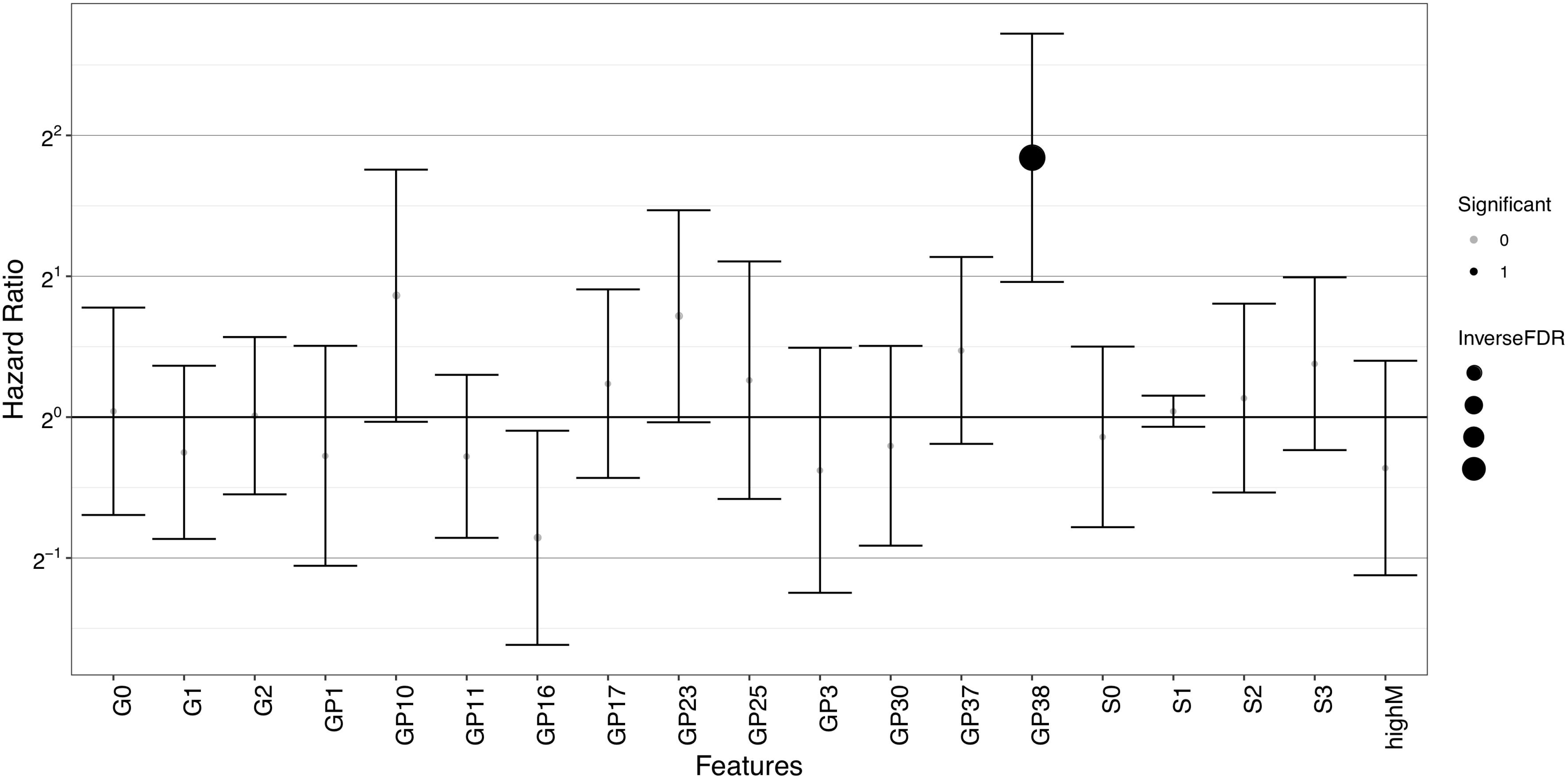
Results of survival analysis (cox-proportional hazard regression) with correction for patient age at diagnosis and tumour infiltrating lymphocyte status (TILs). Survival analysis was run on the set of differentially expressed N-glycan groups. Only one N-glycan, GP38, was significant after correction for multiple testing. Hazard ratios are displayed on a log2 scale with confidence intervals, x-axis = N-glycan groups and y-axis = log2 hazard ratio.

## Discussion

We here present the CAncer bioMarker Prediction Pipeline (CAMPP), a R-based framework for the standardised analysis of high-throughput biological data. Currently the pipeline can handle the higher level normalization, missing value imputation, filtering and correction of a variety of biological data types. The pipeline supports different types of analysis and will provide the user with graphics to support results - all plots displayed in this publication were generated with CAMPP with no or very minimal editing. CAMPP is implemented in such a way that a user is able to run the pipeline on their local computer on datasets with up to 300-500 samples and 40.000 variables. If the user has larger datasets than this, CAMPP may become slow, as especially the cross-validation step in the elastic-net regression will be heavy, it this case, it is advisable to run the pipeline on a server, allocating several cores.

As CAMPP is by far the only framework for automatization of high-throughput data analysis, we here compare the pipeline to a set of other open source tools, found to have comparable functionalities and user demographics. Supplementary Table 1 shows a summary of the CAncer bioMarker Prediction Pipeline (CAMPP), alongside a selection of other softwares, including; *ArrayAnalysis* (Eijssen, et al., 2013), *biojupies* (Torre, et al., 2018), *biowardrobe* (Vallabh, et al., 2018), *Chipster* (Kallio, et al., 2011), *DEWE* (Lopez-Fernandez, et al., 2019), *ExAtlas* (Sharov, et al., 2015), *HiQuAnT* (Bryan, et al., 2016), *Networkanalyst* (Xia, et al., 2015), *PANDA-view* (Chang, et al., 2018), *RobiNA* (Lohse, et al., 2012), *WebMeV* (Wang, et al., 2017). As seen from this comparison the strengths of CAMPP lie both in (I) the variety of analysis it can perform, (II) that the pipeline is able to handle different types of quantitative biological data, from different platforms and (III) that it is flexible in terms of model co-variates. Also, unlike most of the other tools, CAMPP employs *limma* for DE analysis. One advantage of *limma* is that this software has been shown to perform well even with very small sample sizes (Murie, et al., 2009; Seyednasrollah, et al., 2015; Soneson and Delorenzi, 2013), a low power scenario which is not unfamiliar in biomedical research. Lastly, the pipeline is relatively fast to run even with larger gene expression datasets and somewhat robust to different operating systems as it relies on R/Rstudio which are continuously updated to follow system updates.

It should be noted that CAMPP does not support non-aggregated raw array data or raw sequencing reads, and although this was never the scope of the pipeline, which is intended for downstream data analysis, some users may need/prefer a software which can do both at once.

In summary, we here present the CAncer bioMarker Prediction Pipeline (CAMPP) an R-based command line tool for analysis of high-throughput biological data. CAMPP was developed with the intent to provide biomedical researchers with an automated way of “screening” for biomolecules of interest, while ensuring a standardised framework for data normalization and statistical analysis.

## Acknowledgements

The authors would like to thank Vendela Rissler and Emiliano Maiani for testing CAMPP and for helping us with debugging. Our research is founded by Innovationsfonden (5189-00052B), Danmarks Grundforskningsfond (DNRF125), a LEO foundation exploratory grant (LF17006), Carlsberg Foundation Distinguished Fellowship (CF18-0314), The Danish Council for Independent Research, Natural Science, Project 1 (102517) to EP group. The calculations described in this paper were performed using the DeiC National Life Science Supercomputer at DTU. The funders had no role in study design, data collection or analysis, decision to publish, or preparation of the manuscript.

---

Supplementary Figure 1: Multidimensional scaling plot showing the result of k-means clustering with k = 2. The two clusters support the presumed difference between N-glycan abundances in normal interstitial fluid vs tumour interstitial fluid samples.

## References

Agarwal, V., et al. Predicting effective microRNA target sites in mammalian mRNAs. Elife 2015;4.

Alcaraz, N., et al. De novo pathway-based biomarker identification. Nucleic Acids Res 2017;45(16):e151.

Berghuis, A.M.S., et al. Detecting Blood-Based Biomarkers in Metastatic Breast Cancer: A Systematic Review of Their Current Status and Clinical Utility. Int J Mol Sci 2017;18(2).

Bolstad, B.M., et al. A comparison of normalization methods for high density oligonucleotide array data based on variance and bias. Bioinformatics 2003;19(2):185–193.

Bryan, K., et al. HiQuant: Rapid Postquantification Analysis of Large-Scale MS-Generated Proteomics Data. J Proteome Res 2016;15(6):2072–2079.

Cancer Genome Atlas, N. Comprehensive molecular portraits of human breast tumours. Nature 2012;490(7418):61–70.

Celton, M., et al. Comparative analysis of missing value imputation methods to improve clustering and interpretation of microarray experiments. BMC Genomics 2010;11:15.

Chang, C., et al. PANDA-view: an easy-to-use tool for statistical analysis and visualization of quantitative proteomics data. Bioinformatics 2018;34(20):3594–3596.

Chou, C.H., et al. miRTarBase update 2018: a resource for experimentally validated microRNA-target interactions. Nucleic Acids Res 2018;46(D1):D296–D302.

D’Amato, V., et al. Mechanisms of lapatinib resistance in HER2-driven breast cancer. Cancer Treat Rev 2015;41(10):877–883.

Dai, X., et al. Cancer Hallmarks, Biomarkers and Breast Cancer Molecular Subtypes. J Cancer 2016;7(10):1281–1294.

Delignette-Muller, M.L. and Christophe, D. fitdistrplus: An R package for fitting distributions. Journal of Statistical Software 2015(64.4):1–34.

Duffy, M.J., et al. Clinical use of biomarkers in breast cancer: Updated guidelines from the European Group on Tumor Markers (EGTM). Eur J Cancer 2017;75:284–298.

Eijssen, L.M., et al. User-friendly solutions for microarray quality control and pre-processing on ArrayAnalysis.org. Nucleic Acids Res 2013;41(Web Server issue):W71–76.

El-Gebali, S., et al. Solute carriers (SLCs) in cancer. Mol Aspects Med 2013;34(2-3):719–734.

Fridman, W.H., et al. The immune contexture in human tumours: impact on clinical outcome. Nat Rev Cancer 2012;12(4):298–306.

Friedman, J., Hastie, T. and Tibshirani, R. Regularization Paths for Generalized Linear Models via Coordinate Descent. J Stat Softw 2010;33(1):1–22.

Gao, T., et al. Transcriptome analysis reveals the effect of oral contraceptive use on cervical cancer. Mol Med Rep 2014;10(4):1703–1708.

Ghosh, D. and Poisson, L.M. "Omics" data and levels of evidence for biomarker discovery. Genomics 2009;93(1):13–16.

Habashy, H.O., et al. RERG (Ras-like, oestrogen-regulated, growth-inhibitor) expression in breast cancer: a marker of ER− positive luminal-like subtype. Breast Cancer Res Treat 2011;128(2):315–326.

Hansen, T.V., et al. High-density SNP arrays improve detection of HER2 amplification and polyploidy in breast tumors. BMC Cancer 2015;15:35.

Hastie, T., et al. Impute: Imputation for microarray data. 2018(R package version 1.56.0.).

Huang, H.H., Liu, X.Y. and Liang, Y. Feature Selection and Cancer Classification via Sparse Logistic Regression with the Hybrid L1/2 +2 Regularization. PLoS One 2016;11(5):e0149675.

Ioannidis, J.P., et al. Repeatability of published microarray gene expression analyses. Nat Genet 2009;41(2):149–155.

Jabeen, S., et al. Noninvasive profiling of serum cytokines in breast cancer patients and clinicopathological characteristics. Oncoimmunology 2019;8(2):e1537691.

Jensen, L.J., et al. STRING 8--a global view on proteins and their functional interactions in 630 organisms. Nucleic Acids Res 2009;37(Database issue):D412–416.

Kallio, M.A., et al. Chipster: user-friendly analysis software for microarray and other high-throughput data. BMC Genomics 2011;12:507.

Kammers, K., et al. Detecting Significant Changes in Protein Abundance. EuPA Open Proteom 2015;7:11–19.

Kern, S.E. Why your new cancer biomarker may never work: recurrent patterns and remarkable diversity in biomarker failures. Cancer Res 2012;72(23):6097–6101.

Khan, J., Lieberman, J.A. and Lockwood, C.M. Variability in, variability out: best practice recommendations to standardize pre-analytical variables in the detection of circulating and tissue microRNAs. Clin Chem Lab Med 2017;55(5):608–621.

Kursa, M.B. Robustness of Random Forest-based gene selection methods. BMC Bioinformatics 2014;15:8.

La Thangue, N.B. and Kerr, D.J. Predictive biomarkers: a paradigm shift towards personalized cancer medicine. Nat Rev Clin Oncol 2011;8(10):587–596.

Langfelder, P. and Horvath, S. WGCNA: an R package for weighted correlation network analysis. BMC Bioinformatics 2008;9:559.

Leek, J.T. and Storey, J.D. Capturing heterogeneity in gene expression studies by surrogate variable analysis. PLoS Genet 2007;3(9):1724–1735.

List, M., Ebert, P. and Albrecht, F. Ten Simple Rules for Developing Usable Software in Computational Biology. PLoS Comput Biol 2017;13(1):e1005265.

Liu, J., et al. An Integrated TCGA Pan-Cancer Clinical Data Resource to Drive High-Quality Survival Outcome Analytics. Cell 2018;173(2):400–416 e411.

Lo, R., et al. Estrogen receptor-dependent regulation of CYP2B6 in human breast cancer cells. Biochim Biophys Acta 2010;1799(5-6):469–479.

Lohse, M., et al. RobiNA: a user-friendly, integrated software solution for RNA-Seq-based transcriptomics. Nucleic Acids Res 2012;40(Web Server issue):W622–627.

Lopez-Fernandez, H., et al. DEWE: A novel tool for executing differential expression RNA-Seq workflows in biomedical research. Comput Biol Med 2019;107:197–205.

Luo, J., et al. A comparison of batch effect removal methods for enhancement of prediction performance using MAQC-II microarray gene expression data. Pharmacogenomics J 2010;10(4):278–291.

Malta, T.M., et al. Machine Learning Identifies Stemness Features Associated with Oncogenic Dedifferentiation. Cell 2018;173(2):338–354 e315.

McDermott, J.E., et al. Challenges in Biomarker Discovery: Combining Expert Insights with Statistical Analysis of Complex Omics Data. Expert Opin Med Diagn 2013;7(1):37–51.

Merrick, B.A., et al. Platforms for biomarker analysis using high-throughput approaches in genomics, transcriptomics, proteomics, metabolomics, and bioinformatics. IARC Sci Publ 2011(163):121–142.

Minciacchi, V.R., et al. Extracellular vesicles for liquid biopsy in prostate cancer: where are we and where are we headed? Prostate Cancer Prostatic Dis 2017;20(3):251–258.

Morris, C.N. Parametric empirical Bayes inference: theory and applications. Journal of the American statistical Association 1983;78(381):47–55.

Murie, C., et al. Comparison of small n statistical tests of differential expression applied to microarrays. BMC Bioinformatics 2009;10:45.

Naderi, A. C1orf64 is a novel androgen receptor target gene and coregulator that interacts with 14-3-3 protein in breast cancer. Oncotarget 2017;8(34):57907–57933.

Nicolle, R., et al. Prognostic Biomarkers in Pancreatic Cancer: Avoiding Errata When Using the TCGA Dataset. Cancers (Basel) 2019;11(1).

Paculova, H. and Kohoutek, J. The emerging roles of CDK12 in tumorigenesis. Cell Div 2017;12:7.

Papaleo, E., Gromova, I. and Gromov, P. Gaining insights into cancer biology through exploration of the cancer secretome using proteomic and bioinformatic tools. Expert Rev Proteomics 2017;14(11):1021–1035.

Parker, J.S., et al. Supervised risk predictor of breast cancer based on intrinsic subtypes. J Clin Oncol 2009;27(8):1160–1167.

Plautz, G.E., Modi, A. and Wang, L.X. ERBB2 amplicon passenger genes: A novel class of breast cancer antigens. Cancer Res 2014:2897–2897.

Qin, L.X. and Levine, D.A. Study design and data analysis considerations for the discovery of prognostic molecular biomarkers: a case study of progression free survival in advanced serous ovarian cancer. BMC Med Genomics 2016;9(1):27.

Quackenbush, J. Microarray data normalization and transformation. Nat Genet 2002;32 Suppl:496–501.

Rakha, E.A., et al. Biologic and clinical characteristics of breast cancer with single hormone receptor positive phenotype. J Clin Oncol 2007;25(30):4772–4778.

Ritchie, M.E., et al. limma powers differential expression analyses for RNA-sequencing and microarray studies. Nucleic Acids Res 2015;43(7):e47.

Robin, X., et al. pROC: an open-source package for R and S+ to analyze and compare ROC curves. BMC Bioinformatics 2011;12:77.

Robinson, M.D. and Oshlack, A. A scaling normalization method for differential expression analysis of RNA-seq data. Genome Biol 2010;11(3):R25.

Ru, Y., et al. The multiMiR R package and database: integration of microRNA-target interactions along with their disease and drug associations. Nucleic Acids Res 2014;42(17):e133.

Schroder, M.S., et al. survcomp: an R/Bioconductor package for performance assessment and comparison of survival models. Bioinformatics 2011;27(22):3206–3208.

Scrucca, L., et al. mclust 5: Clustering, Classification and Density Estimation Using Gaussian Finite Mixture Models. R J 2016;8(1):289–317.

Seyednasrollah, F., Laiho, A. and Elo, L.L. Comparison of software packages for detecting differential expression in RNA-seq studies. Brief Bioinform 2015;16(1):59–70.

Shannon, P., et al. Cytoscape: a software environment for integrated models of biomolecular interaction networks. Genome Res 2003;13(11):2498–2504.

Sharov, A.A., Schlessinger, D. and Ko, M.S. ExAtlas: An interactive online tool for meta-analysis of gene expression data. J Bioinform Comput Biol 2015;13(6):1550019.

Siu, L.L., et al. Facilitating a culture of responsible and effective sharing of cancer genome data. Nat Med 2016;22(5):464–471.

Soneson, C. and Delorenzi, M. A comparison of methods for differential expression analysis of RNA-seq data. BMC Bioinformatics 2013;14:91.

Stacklies, W., et al. pcaMethods--a bioconductor package providing PCA methods for incomplete data. Bioinformatics 2007;23(9):1164–1167.

Su, D., et al. Role of ERRF, a novel ER-related nuclear factor, in the growth control of ER-positive human breast cancer cells. Am J Pathol 2012;180(3):1189–1201.

Swan, A.L., et al. Application of machine learning to proteomics data: classification and biomarker identification in postgenomics biology. OMICS 2013;17(12):595–610.

Tang, Z.M., et al. Serum tumor-associated autoantibodies as diagnostic biomarkers for lung cancer: A systematic review and meta-analysis. PLoS One 2017;12(7):e0182117.

Terkelsen, T., et al. N-glycan signatures identified in tumor interstitial fluid and serum of breast cancer patients: association with tumor biology and clinical outcome. Mol Oncol 2018;12(6):972–990.

Tiberio, P., et al. Challenges in using circulating miRNAs as cancer biomarkers. Biomed Res Int 2015;2015:731479.

Torre, D., Lachmann, A. and Ma’ayan, A. BioJupies: Automated Generation of Interactive Notebooks for RNA-Seq Data Analysis in the Cloud. Cell Syst 2018;7(5):556–561 e553.

Uttley, L., et al. Building the Evidence Base of Blood-Based Biomarkers for Early Detection of Cancer: A Rapid Systematic Mapping Review. EBioMedicine 2016;10:164–173.

Vallabh, S., Kartashov, A.V. and Barski, A. Analysis of ChIP-Seq and RNA-Seq Data with BioWardrobe. Methods Mol Biol 2018;1783:343–360.

van Ooijen, M.P., et al. Identification of differentially expressed peptides in high-throughput proteomics data. Brief Bioinform 2018;19(5):971–981.

Vieira, A.F. and Schmitt, F. An Update on Breast Cancer Multigene Prognostic Tests-Emergent Clinical Biomarkers. Front Med (Lausanne) 2018;5:248.

Wang, E., et al. Disease Biomarkers for Precision Medicine: Challenges and Future Opportunities. Genomics Proteomics Bioinformatics 2017;15(2):57–58.

Wang, Y.E., et al. WebMeV: A Cloud Platform for Analyzing and Visualizing Cancer Genomic Data. Cancer Res 2017;77(21):e11–e14.

Witwer, K.W. Circulating microRNA biomarker studies: pitfalls and potential solutions. Clin Chem 2015;61(1):56–63.

Xia, J., Gill, E.E. and Hancock, R.E. NetworkAnalyst for statistical, visual and network-based meta-analysis of gene expression data. Nat Protoc 2015;10(6):823–844.

Yen, M.C., et al. Solute Carrier Family 27 Member 4 (SLC27A4) Enhances Cell Growth, Migration, and Invasion in Breast Cancer Cells. Int J Mol Sci 2018;19(11).

Yotsukura, S. and Mamitsuka, H. Evaluation of serum-based cancer biomarkers: a brief review from a clinical and computational viewpoint. Crit Rev Oncol Hematol 2015;93(2):103–115.

