## Supplementary figures and images for "CAncer bioMarker Prediction Pipeline (CAMPP) - A standardised and user-friendly framework for the analysis of quantitative biological data"

### Supplementary Figure S1

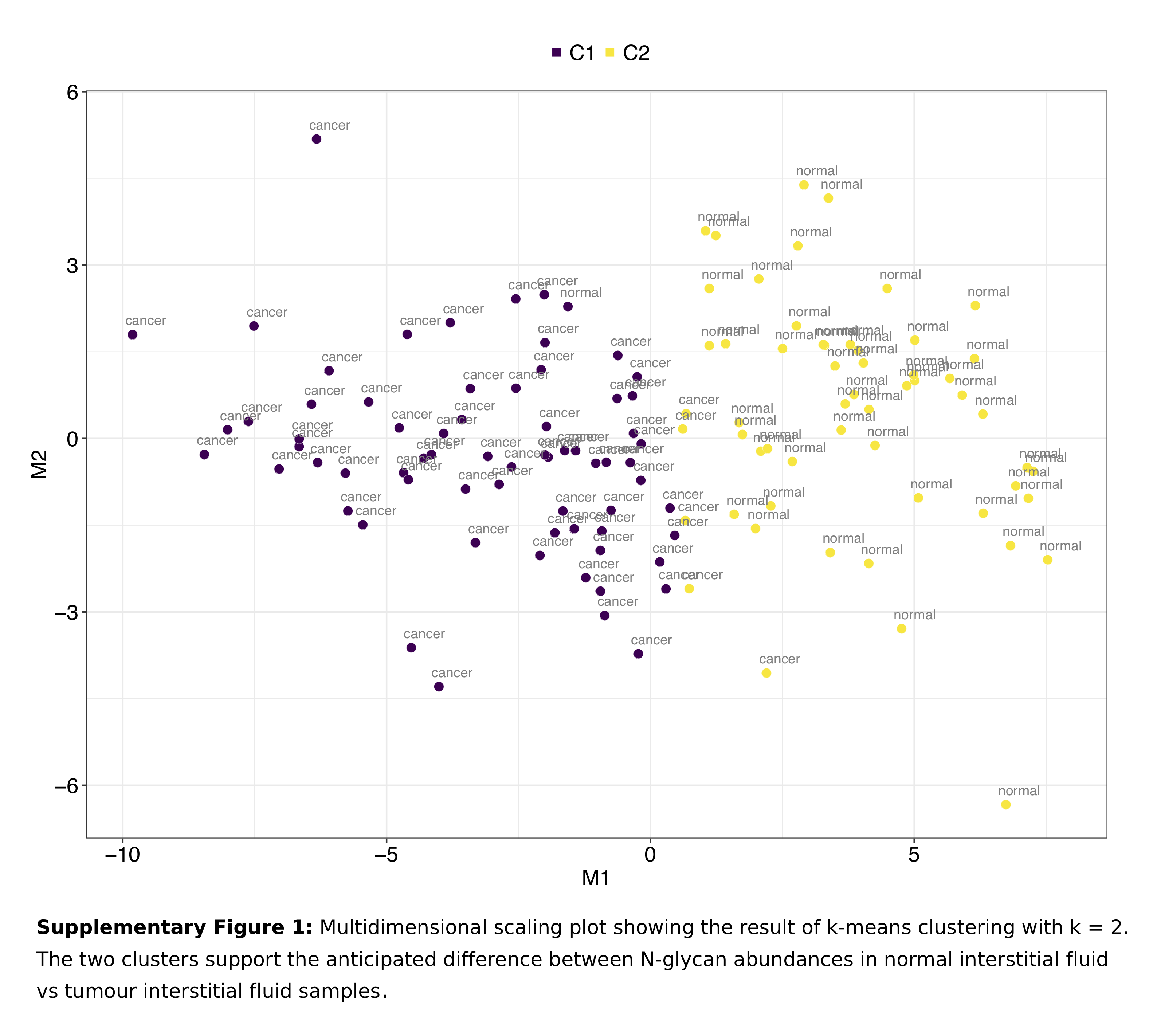
