## Supplementary TableS1 for "CAncer bioMarker Prediction Pipeline (CAMPP) - A standardised and user-friendly framework for the analysis of quantitative biological data"

Table 2 - Comparison of the CAncer bioMarker Prediction Pipeline (CAMPP) with other tools for high-throughput data analysis.

| | Data Type | Platform | Pre-preprocessing and Analyses | Graphics/Plots | Format | Software |
|---|---|---|---|---|---|---|
| **Array Analysis** | Gene expression data | Array (Affymetrix, Illumina) | Quality control (QC) and pre-processing | QC plots, Barplots, Heatmaps, Density plots, Dendo | Graphical web interface | http://www.arrayanalysis.org/ (Eijssen, et al., 2013) |
| | | | Alignment of reads to reference genome | | | |
| | | | Quantification and merging into matrix | | | |
| | | | Normalization/Tranformation | | | |
| | | | Differential expression analysis | Histogram, Boxplots, Barplots | | |
| | | | Pathway enrichment analysis | Pathway graphics plots | | |
| **biojupies** | Gene expression data | Next-Generation Sequencing | Quality control (QC) and pre-processing | | Graphical web interface | https://amp.pharm.mssm.edu/biojupies/ (Torre, et al., 2018) |
| | | | Alignment of reads to reference genome | | | |
| | | | Quantification and merging into matrix | | | |
| | | | Normalization/Tranformation | | | |
| | | | Principal Component Analysis / t-distributed stochastic neighbor embedding | PCA/t-sne plots (3D) | | |
| | | | Hierarchical clustering | Heatmaps | | |
| | | | Differential expression analysis | Volcano plots, Heatmaps, MA plots | | |
| | | | Gene ontology enrichment / Transcription factor and kinase enrichment | Barplots, Dotplots | | |
| | | | Gene pathway enrichment analysis | Pathway graphics plots | | |
| | | | MiRNA enrichment analysis | Barplots, Dotplots | | |
| | | | Identification of small molecules to mimic or reverse DE gene signatures | | | |
| **biowardrobe** | Gene expression data | Next-Generation Sequencing | Quality control (QC) and pre-processing | QC plots, Barplots, Boxplots, Density plots | Download, Install, Graphical interface | https://biowardrobe.com/landing (Vallabh, et al., 2018) |
| | ChIP-sequencing data | | Alignment of reads to reference genome | Pie charts, line graphs | | |
| | | | Quantification and merging into matrix | | | |
| | | | Normalization/Tranformation | | | |
| | | | Principal Component Analysis | | | |
| | | | Identification of islands of enrichment for ChIPs and assignment to a gene region | | | |
| | | | Irreproducible Discovery Rate (IDR) measure/statistic | Sample rank vs IDR scatterplots | | |
| | | | Differential gene expression analysis | Density plots, Boxplots, Heatmaps | | |
| | | | Differential binding analysis | Density plots, Boxplots, Heatmaps | | |
| **Chipster** | Gene expression data | Array (Affymetrix, Illumina) | Quality control (QC) and pre-processing | QC plots, Barplots, Boxplots, Density plots | Graphical web interface | https://chipster.csc.fi/ (Kallio, et al., 2011) |
| | | Next-Generation Sequencing | Alignment of reads to reference genome | | | |
| | | | Quantification and merging into matrix | Histograms, Idiograms | | |
| | | | Normalization/Tranformation | | | |
| | | | Experiment level quality control | Barplots | | |
| | | | Copy number variation (CNV) analysis | | | |
| | | | Correlation analysis | Correlograms | | |
| | | | T-test / ANOVA | Boxplots | | |
| | | | Differential expression analysis / Survival Analysis | Volcano plots, Heatmaps, Chromosomal position | | |
| | | | Hierarchical/K-means clustering | Dendograms | | |
| | | | Principal Component Analysis | PCA plots | | |
| | | | Gene ontology enrichment analysis | | | |
| | | | Pathway enrichment analysis | Pathway graphics plots | | |
| | | | GO enrichment for miRNA targets | | | |
| **DEWE** | Gene expression data | Next-Generation Sequencing | Quality control (QC) and pre-processing | | Download, Install, Graphical interface | http://www.sing-group.org/dewe/ (Lopez-Fernandez, et al. 2019) |
| | | | Alignment of reads to reference genome | | | http://software.broadinstitute.org/software/igv/ |
| | | | Quantification and merging into matrix | | | |
| | | | Normalization/Tranformation | | | |
| | | | Principal Component Analysis | PCA plots | | |
| | | | Differential expression analysis | Volcano plots, Density plots, Boxplots, Heatmaps, Venn diagram | | |
| | | | Pathway enrichment analysis | Pathway graphics plots | | |
| **Exatlas** | Gene expression data | Array (Affymetrix, Illumina) | Standard meta-analysis | | Graphical web interface | https://lgsun.irp.nia.nih.gov/exatlas/ (Sharov, et al., 2015) |
| | | Next-Generation Sequencing | Correlation Analysis | Tile plots | | |
| | | | Gene set enrichment | Heatmaps, Barplots | | |
| | | | Analysis of Variance (ANOVA) | Heatmaps, Barplots | | |
| | | | Principal Component Analysis | PCA plots | | |
| **HiQuAnT** | Quantitative Proteomics | Mass spectrometry data | Missing value imputation | | Download, Install, Graphical interface | http://hiquant.primesdb.eu/ (Bryan, et al., 2016) |
| | | | Normalization/Tranformation | Boxplots | | |
| | | | Differential abundance analysis | Heatmaps | | |
| | | | Protein-Protein Interaction Networks | Network graphics plots | | |
| **Networkanalyst** | Gene expression data | Next-Generation Sequencing | Quality control (QC) and pre-processing | Barplots | Graphical web interface | https://www.networkanalyst.ca/ (Xia, et al., 2015) |
| | | | Alignment of reads to reference genome | | | |
| | | | Quantification and merging into matrix | | | |
| | | | Normalization/Tranformation | | | |
| | | | Data Density and Mean-Variance | Diagnostic plots (scatter, density) | | |
| | | | Differential expression analysis | Heatmaps, Volcano plots | | |
| | | | Principal Component Analysis | PCA plots | | |
| | | | Protein-Protein / TF-miRNA 2D/3D Networks | 2D/3D network plots | | |
| | | | Gene-Chemical, Gene-Drug, Gene-Disease 2D/3D Networks | 2D/3D network plots | | |
| **PANDA-view** | Quantitative Proteomics | Mass spectrometry data | Missing value imputation | Boxplots | Download, Install, Graphical interface | https://sourceforge.net/projects/panda-view/ (Chang, et al., 2018) |
| | | | Normalization/Tranformation | | | |
| | | | Fisher exact test | | | |
| | | | T-test / ANOVA | | | |
| | | | Rank sum test | | | |
| | | | Permutation test | | | |
| | | | Differential abundance analysis | Volcano plots, Density plots, Boxplots, Heatmaps | | |
| | | | K-means / Hierarchical clustering | Scaling plots (Scatter plots) | | |
| | | | Principal Component Analysis | Scree plots, Biplots, PCA plots, Prediction plots | | |
| **RobiNA** | Gene expression data | Next-Generation Sequencing | Quality control (QC) and pre-processing | QC plots (lineplot, barplot, histograms, boxplots) | Download, Install, Graphical interface | https://omictools.com/robina-tool (Lohse, et al., 2012) |
| | | | Alignment of reads to reference genome | | | |
| | | | Quantification and merging into matrix | | | |
| | | | Normalization/Tranformation | Diagnostic plots (scatter, density) | | |
| | | | Principal Component Analysis | PCA plots, Dendograms | | |
| | | | Differential expression analysis | Heatmaps | | |
| **WebMeV** | Gene expression data | Next-Generation Sequencing | Normalization/Tranformation | | Graphical web interface | http://mev.tm4.org/ (Wang, et al., 2017) |
| | | | Hierarchical clustering | Heatmaps | | |
| | | | Principal Component Analysis | PCA plots | | |
| | | | Differential expression analysis | Heatmaps, Dotplots | | |
| **CAMPP** | Gene expression data | Array (Affymetrix, Illumina) | Missing value imputation | | Download, Command line tool | https://github.com/ELELAB/CAncer-bioMarker-Prediction-Pipeline-CAMPP |
| | MiRNA expression data | Next-Generation Sequencing | Normalization/Tranformation | | | |
| | Quantitative Proteomics | Mass spectrometry data | Distributional Checks | Scatter plots, Cullen and Frey graph | | |
| | Quantitative Glycomics/lipi | Liquid chromatography mass spect | K-means Clustering | MDS plots | | |
| | | | Multidimensional Scaling Analysis | MDS plots | | |
| | | | Differential Expression/Abundance Analysis | Heatmaps | | |
| | | | LASSO/Elastic-Net Regression | Barplots | | |
| | | | Survival Analysis | Stemplots | | |
| | | | Weighed Gene Co-expression Network Analysis | Dendograms, heatmaps | | |
| | | | Correlation Analysis | Dotplots, Correlograms | | |
| | | | Protein-Protein and Gene-MiRNA Interaction Networks | Archdigram | | |
